# SFI1 and centrin form a distal end complex critical for proper centriole architecture and ciliogenesis

**DOI:** 10.1101/2021.10.05.463184

**Authors:** Imène B. Bouhlel, Marine. H. Laporte, Eloïse Bertiaux, Alexia Giroud, Susanne Borgers, Juliette Azimzadeh, Michel Bornens, Paul Guichard, Anne Paoletti, Virginie Hamel

## Abstract

Over the course of evolution, the function of the centrosome has been conserved in most eukaryotes, but its core architecture has evolved differently in some clades, as illustrated by the presence of centrioles in humans and a spindle pole body in yeast (SPB). Consistently, the composition of these two core elements has diverged greatly, with the exception of centrin, a protein known to form a complex with Sfi1 in yeast to structurally initiate SPB duplication. Even though SFI1 has been localized to human centrosomes, whether this complex exists at centrioles and whether its function has been conserved is still unclear. Here, using conventional fluorescence and super-resolution microscopies, we demonstrate that human SFI1 is a *bona fide* centriolar protein localizing to the very distal end of the centriole, where it associates with a pool of distal centrin. We also found that both proteins are recruited early during procentriole assembly and that depletion of SFI1 results in the specific loss of the distal pool of centrin, without altering centriole duplication in human cells, in contrast to its function for SPB. Instead, we found that SFI1/centrin complexes are essential for correct centriolar architecture as well as for ciliogenesis. We propose that SFI1/centrin complexes may guide centriole growth to ensure centriole integrity and function as a basal body.

## Introduction

Centrosomes are membrane-less organelles, originally discovered by Theodor Boveri over hundred years ago, which display essential functions in cell biological processes such as cell division (Bornens, 2012; Boveri, 1900). In this case, centrosomes function as the main microtubule nucleating center of the cell (MTOC), forming the two poles of the mitotic spindle that segregates equally the genetic material into the two daughter cells.

While the centrosome is conserved in functional terms in almost all higher eukaryotes-seed plants excepted-, its structure, revealed by numerous electron microscopy studies, has diverged throughout evolution in some species (Azimzadeh, 2014; Ito and Bettencourt-Dias, 2018). In most higher eukaryotes, such as mammals, the centrosome is a proteinaceous condensate surrounding two highly sophisticated core elements called centrioles. Centrioles are 450 nm long cylindrical structures made of nine microtubule triplets, which duplicate in a conservative manner once per cell cycle during S phase (Azimzadeh and Marshall, 2010). In some species, such as yeast or *Dictyostelium*, centrioles have been lost through evolution and replaced by smaller protein assemblies that retained duplication and microtubule nucleation capabilities (Azimzadeh, 2014; Ito and Bettencourt-Dias, 2018; Nabais et al., 2020). In yeast, the centrosome is called the spindle pole body (SPB) and is composed of a core element made of outer and inner plaques associated with a side-appendage, the half-bridge, controlling its duplication (Kilmartin, 2014; Seybold and Schiebel, 2013).

In agreement with the large structural diversity of centrosomes, the proteins constituting their core elements have also diverged greatly (Carvalho-Santos et al., 2011; Hodges et al., 2010; Ito and Bettencourt-Dias, 2018; Nabais et al., 2020). As an illustration, the evolutionary conserved proteins SAS-6, SAS-4/CPAP, CEP135/Bld10p and POC1, all critical for centriole duplication and assembly are absent in yeast (Carvalho-Santos et al., 2011). More generally, even though some centrosome proteins have been conserved between these two species, only centrins have been clearly characterized as present in both, centrioles and yeast SPBs. In mammals, four centrins, centrin-1 to centrin-4 have been identified (Bauer et al., 2016; Gavet et al., 2003; Middendorp et al., 1997; Salisbury et al., 1984), with centrin-1 expressed in the testis and in the retina (Hart et al., 1999; Wolfrum and Salisbury, 1998) and centrin-4 in ciliated cells (Gavet et al., 2003). Centrin proteins are recruited early to procentrioles in the distal lumen of centrioles (Laoukili et al., 2000; Middendorp et al., 2000; Paoletti et al., 1996). Ultrastructure Expansion Microscopy (U-ExM), amenable to nanoscale protein mapping (Gambarotto et al., 2019), further revealed a dual localization for centrin at the central core region and the very distal end of the centriole (Le Guennec et al., 2020; Steib et al., 2020). Functionally, centrins do not seem to be involved in centrosome duplication (Dantas et al., 2011; Strnad et al., 2007), but are rather required for ciliogenesis and the regulation of cell division regulation (Dantas et al., 2011; Delaval et al., 2011; Prosser and Morrison, 2015).

Budding or fission yeasts contain only a single centrin homolog, named Cdc31. In contrast to centrins, Cdc31 is known to be important for SPB duplication by associating with the protein Sfi1 (Baum et al., 1986; Kilmartin, 2003; Li et al., 2006; Paoletti et al., 2003; Spang et al., 1995; Vallen et al., 1994), an extended α-helix that possess multiple Cdc31-binding domains (Li et al., 2006), and which, upon Cdc31 binding, assembles into a parallel array to form the SPB half-bridge. Assembly of a second array of Sfi1/Cdc31, anti-parallel to the first, and associated with it through Sfi1 C-termini was shown to define the site for daughter SPB assembly, controlling thereby SPB conservative duplication (Bestul et al., 2017; Bouhlel et al., 2015; Lee et al., 2014).

Interestingly, it was shown that SFI1 localises at centrosomes in human cells (Kilmartin, 2003; Kodani et al., 2019) and can interact directly with human centrins *in vitro* (Martinez-Sanz et al., 2010, 2006). However, it remains unclear whether centrins and human SFI1 form a complex at centrioles. Indeed, in contrast to what was found for centrins, it was recently proposed that SFI1 regulates centriole duplication, similar to its function at SPBs, but in this case by stabilizing the proximal protein STIL (Balestra et al., 2013; Kodani et al., 2019). These results raised the possibility that the centrin/SFI1 complex have not been conserved in human centrioles. To test this hypothesis, we have studied further the localization and function of human SFI1 combining cell biology and super-resolution techniques. We first establish that SFI1 is a molecular constituent of the centriole that co-localizes with a distinct pool of centrin2/3 at the very distal tip of centrioles, from early stages of centriole biogenesis in human cells. We further demonstrate that SFI1 is dispensable for early events of centriole duplication but that its depletion leads to the specific loss of the distal pool of centrins and strongly affects centriole architecture as well as ciliogenesis. These results suggest that the conserved SFI1/centrin complex has a major role, not in the early stages of centriole duplication, but during their elongation, to ensure correct centriole assembly, with impact on centriole / basal body functions.

## Results

### Human SFI1 is a *bona fide* centriolar component localizing at the very distal end

Human SFI1 is an evolutionary conserved protein of 1242 amino acids that contains 23 characteristic SFI1 repeats (Kilmartin, 2003; Li et al., 2006) (**Figure S1**). SFI1 has been shown to localize at centrosomes (Kilmartin, 2003; Kodani et al., 2019) as well as at centriolar satellites during S phase (Kodani et al., 2015). To investigate whether SFI1 is a *bona fide* centriolar component, we raised and affinity-purified a polyclonal antibody against a C-terminus fragment of the protein encompassing residues 1021 to 1240 (**Figure S1**). First, immunofluorescence analysis of cycling immortalized hTERT RPE-1 cells (hereafter referred to as RPE-1) co-stained for the centrosomal marker γ-tubulin and SFI1, confirmed its localization at centrosomes throughout the cell cycle (**Figure 1A**). Moreover, using the centrin 20H5 monoclonal antibody, which recognizes human centrin 2 and centrin 3 (Middendorp et al., 1997; Paoletti et al., 1996; Sanders and Salisbury, 1994), we found that SFI1 localizes at centrioles (**Figure 1B**). To further investigate the precise localization of SFI1 at centrioles, we turned to super-resolution ultrastructure expansion microscopy (U-ExM) (Gambarotto et al., 2021, 2019). Interestingly, we found that the C-terminus of SFI1 localizes as a distinct dot at the very distal tip in mature centrioles both in RPE-1 and in osteosarcoma U2OS cells (**Figure 1C-E** and **Figure S2A, J**). To ascertain the specificity of this signal, we analyzed SFI1 distribution in RPE-1 cells depleted of SFI1 upon siRNA treatment, as previously described (Balestra et al., 2013). We found that the distal dot corresponding to SFI1 disappeared, confirming the specificity of the signal (**Figure S2B-D**). The specificity of this localization was further tested using the commercially available SFI1 antibody (13550-1-AP, Proteintech Europe). Consistently, we found the same localization at the distal extremity, which decreases upon siRNA depletion in RPE-1 cells (**Figure S2E-I, K**).

**Figure 1.**
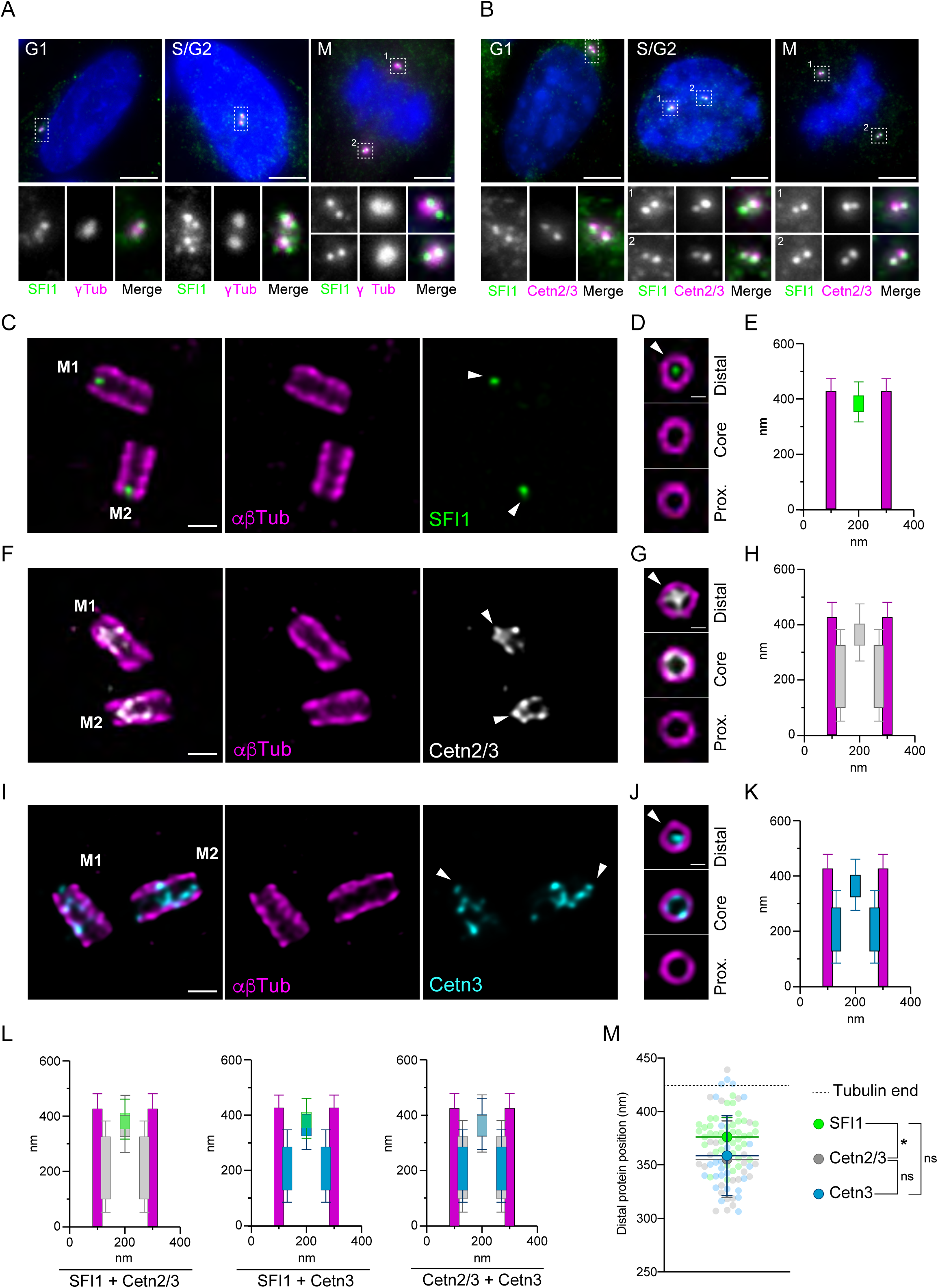
SFI1 is a centriolar protein co-localizing with Centrin 2 and 3 at the distal tip of centrioles. **(A, B)** Representative confocal images of cycling RPE-1 cells stained for SFI1 (green) and γ-Tubulin (magenta) (A) or SFI1 (green) and Centrin 2/3 (magenta) (B). Scale bar= 5µm. Dashed-line squares correspond to insets. (**C, F, I)** Representative confocal images of expanded centrioles from RPE-1 cells stained for α/β-tubulin (αβTub, magenta) and SFI1 (green) (C), Centrin 2/3 (Cetn2/3, grey) (F) or Centrin 3 (Cetn3, cyan) (I). The white arrowhead indicates SFI1 (C) and Centrin (F, I) distal dots at centrioles. Scale bars: 200 nm. **(D, G, J)** Top view images of expanded centrioles from RPE-1 cells stained for α/β-tubulin ((αβTub, magenta) and SFI1 (green) (D), Centrin2/3 (Cetn2/3, grey) (G) or Centrin 3 (Cetn3, cyan) (J) unveiling the distal localization of SFI1 at centrioles. The white arrowhead indicates SFI1 (D) and Centrin (G, J) distal dots at centrioles. Scale bars: 100 nm. **(E, H, K)** Average position of SFI1 (E), Centrin 2/3 (H), Centrin 3 (K) alongside the centriole. (**L**) Superposition of the average position of SFI1 with Centrin2/3 (left), SFI1 with Centrin 3 (middle) and Centrin 2/3 with Centrin 3 (right). **(M)** Position of SFI1 and Centrin signals at the distal centriolar region in nm. (**E, H, K, L, M**) Averages +/- SD: SFI1= 376 +/- 18 nm; Centrin2/3: 355 +/- 36 nm; Centrin3: 359 +/- 37 nm. N = 41, 25, 24 centrioles for SFI1, Centrin 2/3 and Centrin 3 respectively, from 2 independent experiments. One-way ANOVA followed by Bonferroni post-hoc test (SFI1 vs Cetn2/3 p=0.0196, SFI1 vs Cetn3 p=0.0702, Cetn2/3 vs Cetn3 p=0.999).

We also noted a faint dotty proximal signal that decreases upon SFI1 depletion, possibly reflecting a putative additional location for SFI1 (**Figure S2B, C, F** and **H**, red arrowhead).

We next compared the precise distribution of SFI1 and centrin2/3 at centrioles (**Figure 1C-M**). We found that both centrin2/3 and centrin3 localize as a dot at the distal tip of centrioles, with an additional distribution at the central core region as previously reported (Le Guennec et al., 2020) (**Figure 1F-K**). By measuring and comparing the relative position of the distal dots of SFI1 and centrins compared to the microtubule wall defined by the tubulin staining, we found that the average position of SFI1 is only 20 nm apart from that of centrin2/3, with a non-significant difference for centrin-3, indicating that SFI1 and centrin 2/3 co-localize strictly at the distal end of centrioles (**Figure 1L, M**). Based on this nanometric proximity, and the known interaction between centrins and SFI1 in yeast and human *in vitro* (Bouhlel et al., 2015; Li et al., 2006; Martinez-Sanz et al., 2006), we propose that centrin2/3 and SFI1 form a complex at the distal end of the human centriole.

Next, we decided to monitor the recruitment of the SFI1/centrins complex during centriole assembly. As centrins are recruited to procentrioles during early phases of centriole biogenesis (Middendorp et al., 1997; Paoletti et al., 1996), we investigated whether this was also the case for SFI1. Immunofluorescence analysis of RPE-1 cells in S phase, identified using the nuclear PCNA marker (Takasaki et al., 1981), indicated the presence of more than two dots of SFI1 at centrosomes at this stage (**Figure 2A**), compatible with a recruitment of SFI1 at procentrioles. However, the SFI1 signal appears cloudy, reminiscent to the satellites localization previously described (Kodani et al., 2015). Therefore, we next analyzed duplicating centrioles using U-ExM (**Figure 2B**). We found that SFI1 localizes at procentrioles, and, similarly to centrin2/3 and centrin3 alone, is already present at the growing distal tip of centrioles from nascent procentrioles both in RPE-1 and U2OS cells (**Figure 2B** and **Figure S2A, E, J, K**). This result demonstrates that the SFI1/centrins complex is recruited at the onset of centriole biogenesis. Note that since centrin 2/3 and centrin 3 display similar localizations (**Figure 1M**), we will next refer to the generic term centrin from thereafter for the sake of simplicity.

**Figure 2.**
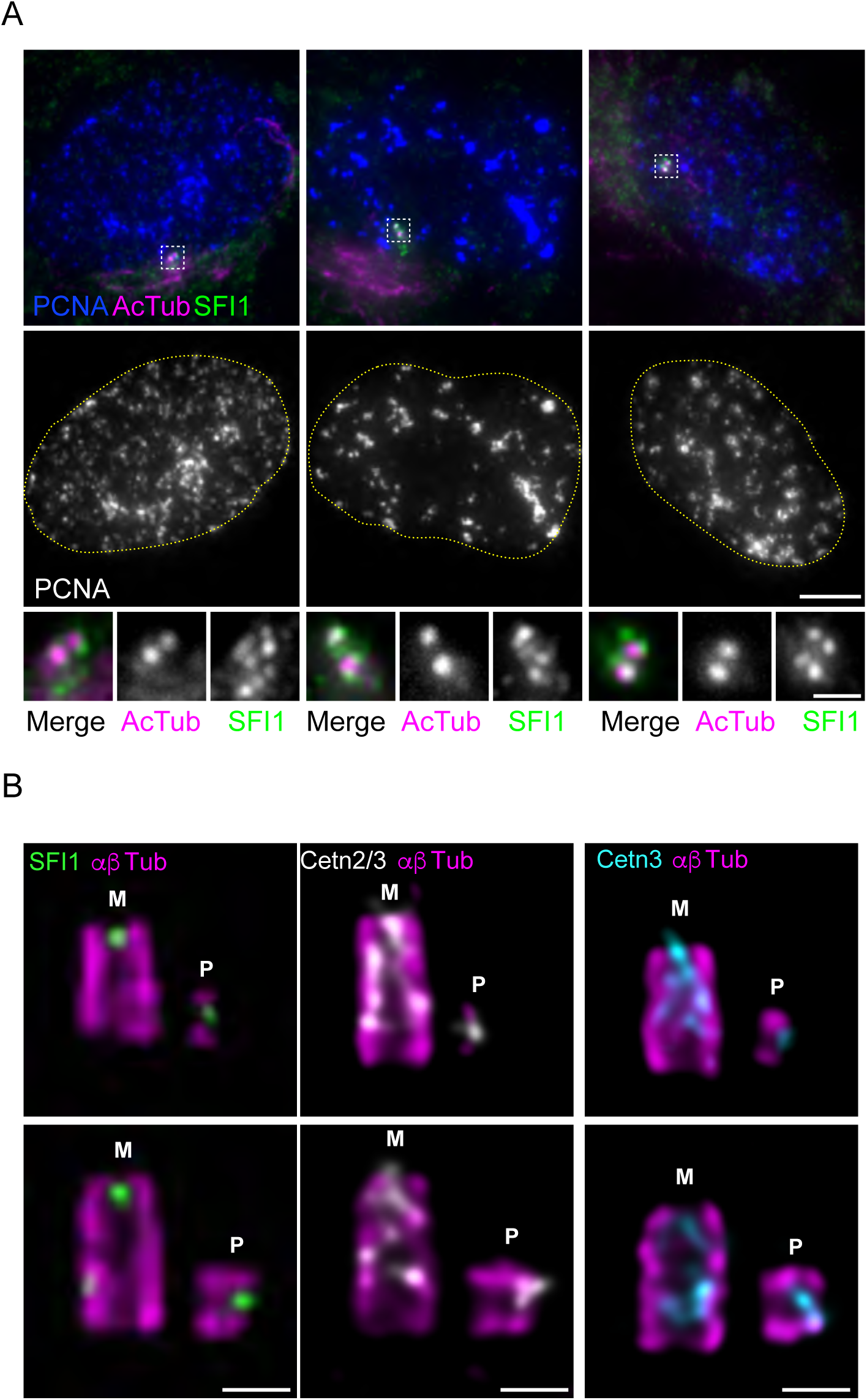
SFI1 and Centrin are recruited at the onset of centriole biogenesis. **(A)** Representative confocal images of RPE-1 cycling cells stained for SFI1 (green), Acetylated Tubuin (AcTub, magenta) and PCNA (blue). DNA boundaries are marked with a yellow dotted line. White dashed line squares correspond to insets. Scale bar= 5 µm. **(B)** Representative confocal images of expanded duplicating centrioles from RPE-1 cells stained for α/β-tubulin (αβTub, magenta) and SFI1 (green, left panel), Centrin 2/3 (Cetn2/3, grey, middle panel) or Centrin 3 (Cetn3, cyan, rigth panel). M stands for mature centriole, P stands for procentriole. Note that both SFI1 and Centrins are recruited very early at procentrioles as a distal dot. Scale bars: 200 nm.

### SFI1 is critical for distal centrin recruitment at centrioles

Next, we assessed the impact of SFI1 depletion on centrin localization at centrioles. To do so, we co-stained control and SFI1-depleted RPE-1 cells with centrin and the distal end protein CP110 as a proxy for centriole’s presence (Schmidt et al., 2009) (**Figure 3A**). We found that centrin signal was strongly reduced upon SFI1 depletion, often solely present at one centriole, while CP110 appeared unchanged (**Figure 3A, B**). To ascertain this observation, we turned again to expansion microscopy. We monitored SFI1 depletion at mature centrioles and found that while most of the control cells were positive for SFI1, 97% of the SFI-depleted cells had lost the centriolar distal dot of SFI1, demonstrating the efficiency of the siRNA treatment (**Figure 3C, D, I**). Similarly, we observed that 97% of centrioles had lost centrins at their distal end (**Figure 3F, G, K**, yellow arrowhead), while keeping intact the centrin pool at the central core region (**Figure 3G**). This result suggests that SFI1 controls specifically the localization of a centrin pool at the distal end of centrioles. To further assess this hypothesis, we depleted the inner scaffold protein POC5, also known to interact with centrin (Azimzadeh et al., 2009) and analyzed the distribution of both centrin and SFI1. Remarkably, we found that the distal pool of centrin remained unchanged while the pool of centrin at the central core region was strongly affected (**Figure 3H, L** and **Figure S2L-P)**. This result demonstrates that centrin forms two distinct complexes, one at the central core relying on the POC5 protein and one at the distal end of centrioles dependent on SFI1. Consistently, SFI1 localization remains unchanged in POC5-depleted cells (**Figure 3E, J** and **Figure S2L-P**), confirming that POC5 specifically affects centrin at the central core region of centrioles and does not impact the distal complex of SFI1/centrin.

**Figure 3.**
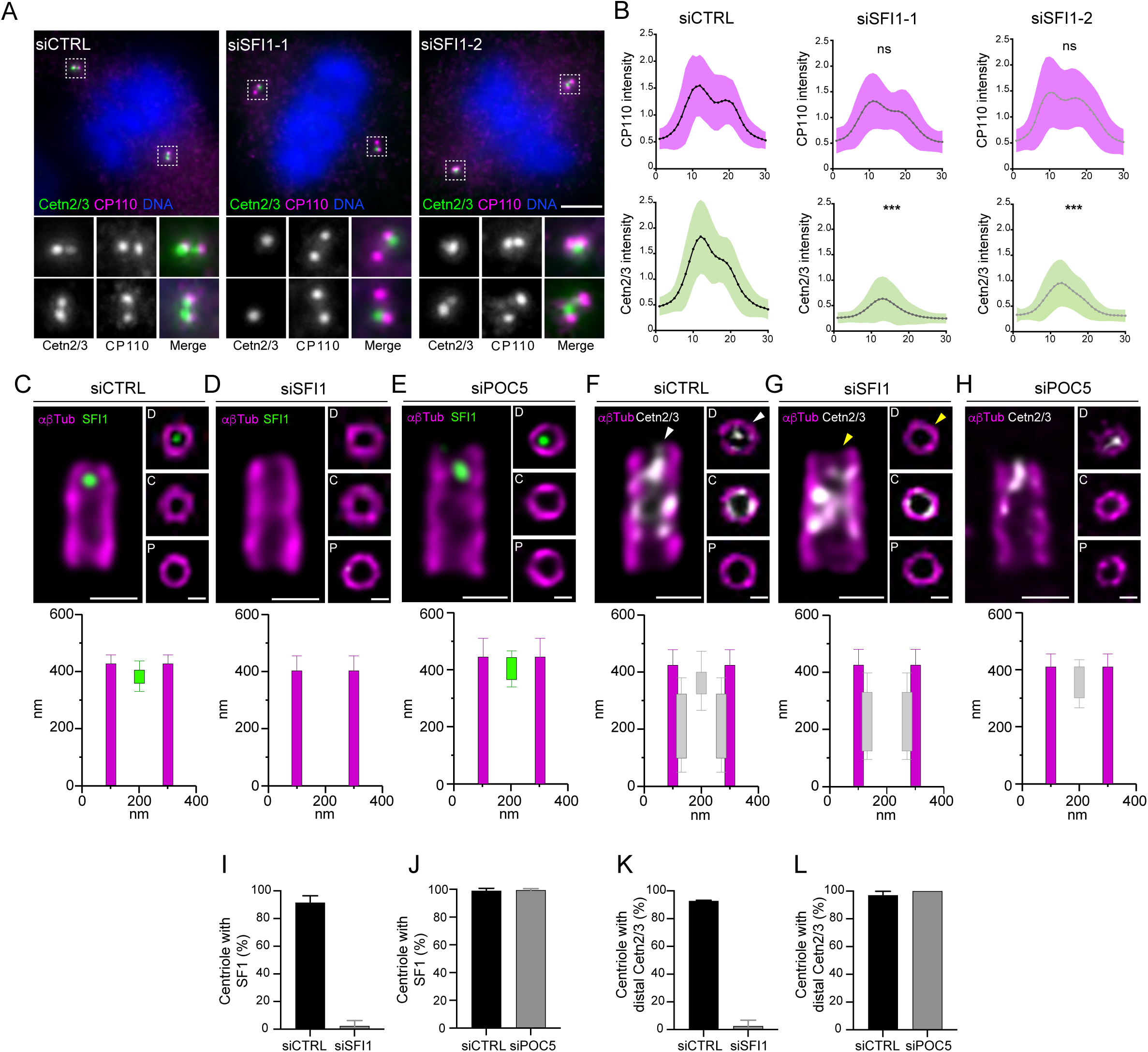
SFI1 depletion prevents distal Centrin recruitment at centrioles. (**A)** Representative confocal images of mitotic control and SFI1-depleted RPE1-1 cells stained for Centrin (Cetn2/3, green) and CP110 (magenta). Scale bar: 5µm. **(B)** CP110 (magenta) and Centrin (green) relative integrated intensities from a plot profile across the 2 centrioles in control and SFI1-depleted cells (siSFI1-1 and siSFI1-2 correspond to two different siRNAs, see material and methods). Averages of AUC (area under the curve) +/- SD are as follows: siControl (CP110) = 1.001+/-0.349, siSFI1-1 (CP110) = 0.9039 +/- 0.279, siSFI1-2 (CP110) = 1.013 +/- 0.345; siControl (Cetn2/3) = 0.994 +/- 0.480, siSFI1-1 (Cetn2/3) = 0.3782 +/- 0.130, siSFI1-2 (Cetn2/3) = 0.548 +/- 0.226. N=60 cells from 3 independent experiments, ***p<0.0001, Student t-test. (**C-E**) Representative confocal images of expanded U2OS centrioles treated with siCTRL (**C**), siSFI1 (**D**) and siPOC5 (**E**) stained for α/β-tubulin (αβTub, magenta) and SFI1 (green). Insets show top views of expanded centrioles at different positions along the centriole (P= proximal, C= central and D= distal). Note that in the absence of SFI1 staining at the distal tip, orientation of the centriole was decided based on the larger diameter in the proximal region compared to the distal one, as previously observed in cryo-tomography (Greenan et al., 2020). Scale bars: 200 nm and 100 nm (inset). Longitudinal and radial localisation of SFI1 in siCTRL (**C**), siSFI1 (**D**) and siPOC5 (**E**) are presented below the corresponding image. Averages +/- SD are as follows: siCTRL: 0 to 427 +/- 31 nm for tubulin and 359 +/- 37 to 405 +/- 37 nm for SFI1, siSFI1: 0 to 403 +/- 52 nm for tubulin, siPOC5: 0 to 445 +/- 66 nm for tubulin and 367 +/- 65 to 443 +/- 68 nm for SFI1. N=25, 40 and 60 centrioles from 3 independent experiments. (**F-H)** Representative confocal image of expanded U2OS centrioles treated with siCTRL (F), siSFI1 (**G**) and siPOC5 (**H**) stained for α/β-tubulin (αβTub, magenta) and Centrin2/3 (Cetn2/3, grey). Insets show top views of expanded centrioles at different positions along the centriole (P= proximal, C= central and D= distal). Note that in the absence of distal centrin staining at the distal tip, orientation of the centriole was decided based on the larger diameter in the proximal region compared to the distal one. Scale bars: 200 nm and 100 nm (inset). White arrowheads points to the distal dot of Centrin that disappears in SFI1-depleted (yellow arrowheads) but not in POC5-depleted centrioles. Longitudinal and radial localization of Centrin in siCTRL (**F**), siSFI1 (**G**) and siPOC5 (**H**) are presented below the corresponding image. Averages +/- SD are as follows: siCTRL : 0 to 424 +/- 55 nm for tubulin and 98 +/- 47 to 391 +/- 39 nm for Centrin, siSFI1 : 0 to 425 +/- 56 nm for tubulin and 122 +/- 26 to 324 +/- 51 nm for Centrin, siPOC5: 0 to 410 +/- 45 nm for tubulin and 301 +/- 49 to 409 +/- 46 nm for Centrin. N=29, 34, 34 centrioles from 3 independent experiments. **(I, J)** Percentage of centrioles containing SFI1 as a distal dot in siSFI (**I**) and siPOC5 (**J**) compared to control cells. Averages +/- SD are as follows: **I**) siCTRL= 91.6% +/- 4.7, siSFI1= 2.3% +/- 4; **J**) siCTRL= 99% +/- 1.7, siPOC5= 99.3% +/- 1.5 N= 3 independent experiments (>80 centrioles per experiment). **(K, L)** Percentage of centriole with a distal Centrin 2/3 signal in siSFI (K) and siPOC5 (L) compared to control cells. Averages +/- SD are as follows: K) siCTRL= 92.7 +/- 0.6, siSFI1= 2.5% +/- 4.3; L) siCTRL= 97% +/- 2, siPOC5= 100% +/- 0, suggesting that POC5 depletion does not impact distal Centrin under these experimental conditions. N= 3 independent experiments (>80 centrioles per experiment).

### SFI1 is important for ciliogenesis, but not for centriole duplication

Since centrin is important for ciliogenesis (Delaval et al., 2011; Prosser and Morrison, 2015), we speculated that this function might be specifically related to the distal SFI1/centrin complex due to its close proximity with the transition zone for cilium formation. Therefore, we first looked at whether SFI1 was still present in ciliogenesis. We found by immunofluorescence and U-ExM that SFI1 localizes and remains at the distal end of the ciliated centrioles in RPE-1 cells (**Figure 4A, B**). Similarly, staining of centrin in those cells revealed that the distal centrin dot also remains in ciliated cells, indicating that the whole complex is retained in these conditions (**Figure 4C**, arrowheads). Next, we investigated the impact of SFI1 depletion on ciliogenesis. As found for centrin depletion (Prosser and Morrison, 2015), we observed that only 26% of SFI1-depleted cell displayed a primary cilium stained with acetylated tubulin whereas ∼75% of control cells displayed were ciliated (**Figure 4D, E**), demonstrating that, as centrin (Prosser and Morrison, 2015), SFI1 is essential for ciliogenesis in human cells.

**Figure 4.**
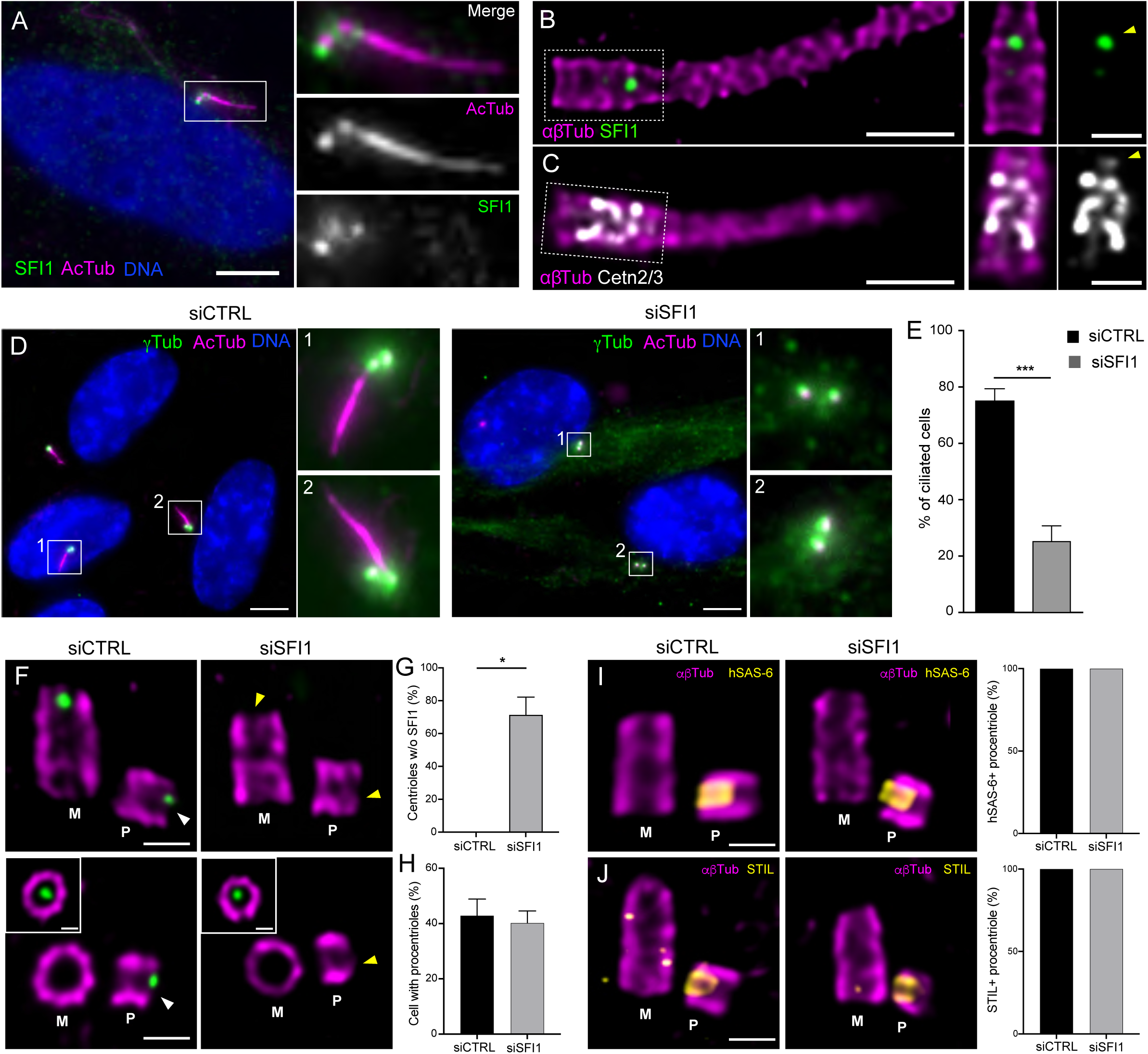
SFI1 is important for ciliogenesis but does not impair centriole duplication. **(A)** Representative confocal images of serum-starved RPE-1 cells stained for SFI1 (green) and acetylated tubulin (AcTub, magenta). Scale bar: 5µm. **(B, C)** Representative confocal images of serum-starved expanded RPE-1 stained for SFI1 (**B**, green) or Centrins (**C**, Cetn2/3, grey) and α/β-tubulin (αβTub, magenta). Insets show single channel depicting the distal localization of SFI1 (**B**, arrowhead) and Centrins (**C**, arrowhead). Scale bars: 500 nm and 200 nm (inset). (**D**) Representative confocal images of serum-starved RPE-1 cells transfected with control or SFI1 siRNA stained for γ-tubulin (γTub, green) and acetylated tubulin (AcTub, magenta) and DNA (DAPI, blue). Scale bar: 5µm. (**E**) Percentage of ciliated cells in the indicated conditions. Averages +/- SD are as follows: siCTRL= 75% +/- 3, siSFI1= 26% +/- 6. N= 3 independent experiments (100 cells per experiment), ***p value=0.0002, unpaired t-test. (**F**) Representative confocal image of expanded duplicating centrioles from siCTRL and siSFI1 U2OS treated cells. Cells were stained for SFI1 (green) and α/β-tubulin (αβTub, magenta). Inset shows a distal position of the mother centriole where SFI1 signal is visible. White arrowhead indicates the position of SFI1 distal dot in the procentriole of control cell, which is lost in SFI1-depleted cells (yellow arrowhead). Scale bars: 200 nm. **(G)** Quantification of the percentage of SFI1-negative procentrioles. Average +/- SD are as follow: siCTRL= 0% +/- 0, siSFI1=71% +/- 11. N = 4 independent experiment (50 centrioles per experiment), *pvalue=0.028, Mann-Whitney test. (**H**) Quantification of the percentage of duplicating centrioles. Averages +/- SD are as follows: siCTRL= 43% +/- 6, siSFI1= 40% +/- 4. N= 7 independent experiment (50 centrioles per experiment), pvalue= 0.3492, Student t-test. (**I**) Representative confocal image of expanded duplicating centrioles from siCTRL and siSFI1 U2OS treated cells. Cells were stained for HsSAS-6 (yellow) and α/β-tubulin (αβTub, magenta). Quantification shows no difference in the percentage of HsSAS-6-positive centrioles in siSFI1-depleted cells compared to control cells. Averages and SD are as follows: siCTRL= 100% +/- 0, siSFI1= 100% +/- 0. N = 3 independent experiments, pvalue>0.999, Mann Whitney test. (**J**) Representative confocal image of expanded duplicating centrioles from siCTRL and siSFI1 U2OS treated cells. Cells were stained for STIL (yellow) and α/β-tubulin (αβTub, magenta). Quantification shows no difference in the percentage of STIL-positive centrioles in siSFI1 compared to control cells. Averages +/- SD are as follows: siCTRL= 100% +/- 0, siSFI1= 100% +/- 0. N = 3 independent experiments, pvalue>0.999, Mann Whitney test.

Next, we wanted to assess the impact of SFI1 depletion on centriole biogenesis as it has been reported to impact centriole duplication (Balestra et al., 2013; Kodani et al., 2019). To do so, we turned to osteosarcoma U2OS cells, widely used to study centriole duplication. As we cannot use centrin as a marker for centriole duplication since SFI1 depletion impairs its localization, we decided to directly monitor the presence of procentrioles using tubulin staining by U-ExM (**Figure 4 F, G**). In contrast to the strong effect on the number of centrin dots (**Figure 3A, B**) (Balestra et al., 2013; Kodani et al., 2019), we found that centriole duplication occurs normally in SFI1-depleted centrioles from U2OS cells (**Figure 4F, G**, yellow arrowheads). We observed no significant difference in the percentage of cells with procentrioles in control or SFI1-depleted cells, with an average of 42% +/-6 and 40% +/-4 of cells harboring procentrioles (**Figure 4H**). To further investigate any putative duplication phenotype, we monitored the presence of the cartwheel proteins HsSAS-6 and STIL at procentrioles, as previous data showed that SFI1-depleted HeLa cells failed to recruit these two proteins to S-phase centrosomes, probably owing to STIL destabilization (Kodani et al., 2019). In contrast, we found that both HsSAS-6 and STIL are recruited in the growing procentrioles of U2OS SFI1-depleted cells (**Figure 4I, J**). Collectively, these data demonstrate that SFI1 depletion does not affect the initiation of procentriole assembly in human U2OS cells under our experimental conditions as opposed to its role in SPB duplication. This unveiled that, despite the conservation of the Centrin/SFI1 complex between yeast and mammals, its function in duplication does not seem to be conserved.

### SFI1 is important for centriole integrity

Since SFI1 depletion leads to ciliogenesis defects, a process linked to the function of the mature centriole, we next wondered whether the architecture of the mature centriole itself could be affected upon SFI1 depletion in cycling U2OS cells using U-ExM (**Figure 5** and **Figure S3**). Strikingly, we observed that SFI1-depleted centrioles had lost the canonical round shape of the microtubule wall, highlighted by the quantification of the roundness index (**Figure 5A, B** and **Video 1**). In contrast, we did not notice any significant difference in overall centriole diameter nor average length even though we observed a wider distribution of sizes with shorter and longer centrioles (**Figure 5C** and **Figure S3A, B**). Furthermore, we found that 35% of centrioles were structurally abnormal in SFI1-depleted cells (**Figure 5D, E, Figure S3** and **Video 1**), with often opened, wider or shorter microtubule walls (**Figure S3**). This structural phenotype suggests that SFI1 is crucial for the integrity of centriole’s architecture, likely explaining the observed phenotype such as the defective ciliogenesis. Interestingly, we noticed that the absence of microtubule wall observed in SFI1-depleted abnormal centrioles was correlated with the lack of the inner scaffold localization of centrin (**Figure 5F, G**), while CP110 was still present at the tip of these centriolar microtubule wall structures (**Figure S3C, D**). As our data showed that SFI1 depletion specifically leads to distal centrin loss, it is likely that the absence of centrin at the central core may be an indirect consequence of the microtubule wall defect.

**Figure 5.**
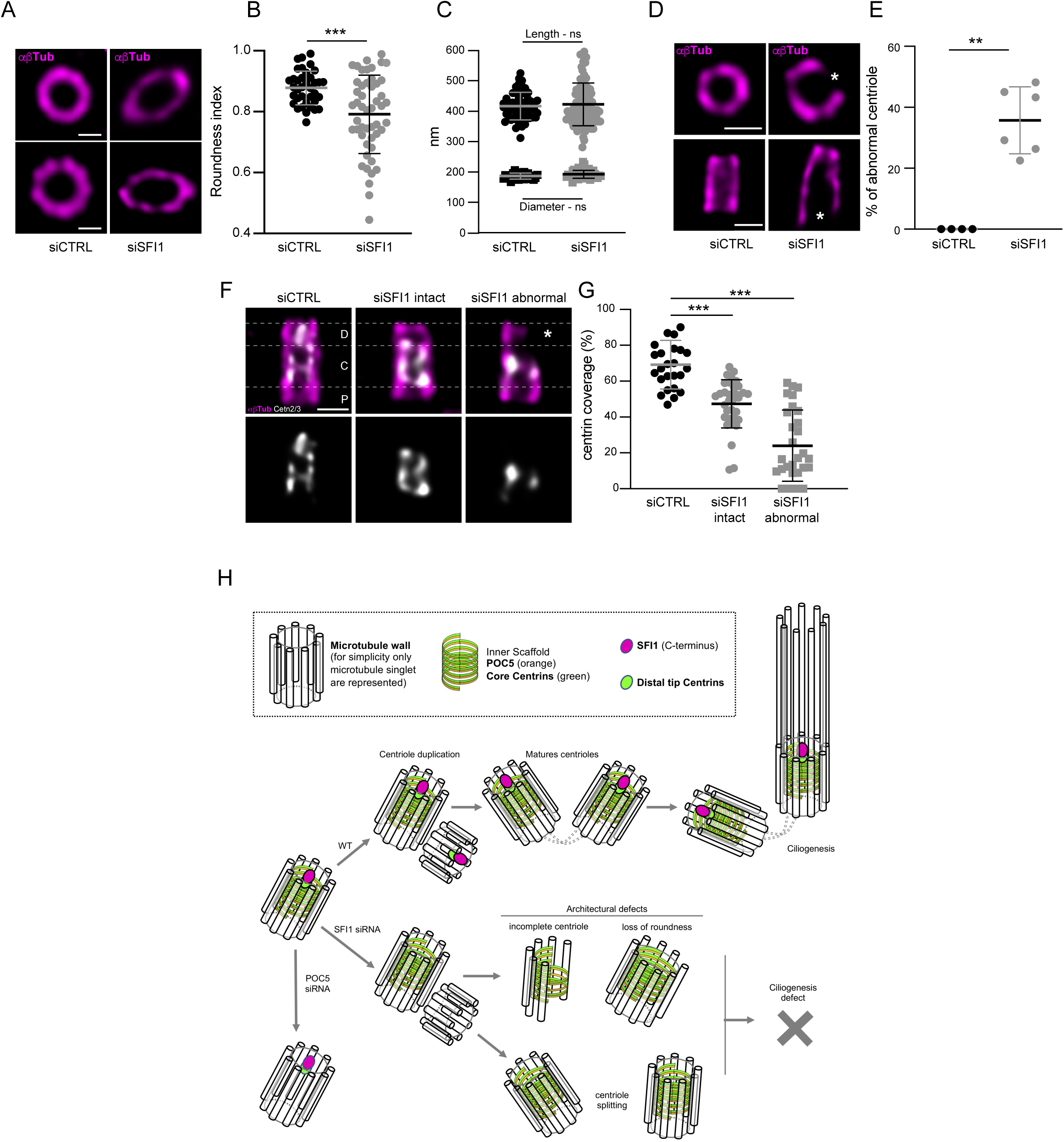
SFI1 depletion leads to centriole structural defects. **(A)** Top views of expanded U2OS centrioles treated with siCTRL or siSFI1 stained for α/β-tubulin (αβTub, magenta). Scale bars: 200 nm **(B)** Roundness index calculated for siCTRL and siSFI1-treated centrioles. Averages +/- SD: siCTRL= 0.88+/-0.05, siSFI1-1= 0.79+/-0.12. N= 37 and 50 for siCTRL and siSFI1 respectively from 4 independent experiments, ***p= 0.0002, unpaired t-test. **(C)** Length (circle) and diameter (square) of expanded centrioles in siCTRL or siSFI1-treated cells. Averages +/- SD are as follows: siCTRL= 417 +/- 45 nm (length) and 188 +/- 10 nm (diameter), siSFI1= 423 +/- 71 nm (length) and 193.5 +/- 13 nm (diameter). N= 50-90 for length and 30-40 for diameter from 4 independent experiments, p=0.8440 (length), p=0.079 (diameter), Mann-Whitney test. **(D)** Representative confocal images of expanded U2OS centrioles from siCTRL and siSFI1-treated cells stained for α/β-tubulin (αβTub, magenta). White stars point to broken microtubule wall. Scale bars: 200 nm. **(E)** Percentage of abnormal centrioles in the indicated conditions. Averages +/- SD are as follows: siCTRL= 0% +/- 0, siSFI1-1= 35.7% +/- 11. N= 4 and 6 independent experiments for siCTRL and siSFI1 respectively, p=0.0095, Mann-Whitney test. (**F**) Representative confocal images of expanded U2OS centrioles from siCTRL and siSFI1-treated cells stained for α/β-tubulin (αβTub, magenta) and Centrin (Cetn2/3, grey). White dashed lines delimitate the proximal, central and distal regions. White star points to the broken microtubule wall. Scale bar: 200 nm. **(G)** Centrin coverage (% of the total tubulin length) along the centriole in the indicated conditions. Averages +/- SD are as follows: siCTRL: 69% +/-13, siSFI1: 47% +/-13, siSFI1 abnormal: 24% +/- 20. N=25, 34, 29 centrioles for siCTRL, siSFI1, siSFI1 abnormal respectively from 2 independent experiments. One-way ANOVA followed by Tukey’s post-hoc test, siCTRL vs siSFI1-1, p<0.0001, siCTRL vs siSFI1-1 abnormal, p<0.0001. (**H**) Model of Centrin/SFI1 localization and function at the distal end of human centrioles.

Overall, these results demonstrated that SFI1 is crucial for the establishment of the correct centriolar architecture. As SFI1 is essential for the localization of centrin at the distal end of centrioles from the early steps of procentriole assembly, it is possible that Centrin and SFI1 acts as a scaffolding complex that would guide proper centriole biogenesis during centriole elongation, thus ensuring centriole integrity that is necessary for mature centriole function (**Figure 5H**).

## Discussion

The evolutionary origin of the centriole remains an enigma, but its near-ubiquitous existence in eukaryotes, as well as phylogenetic analyses, have led to propose that this organelle was already present in the Last Eukaryotic Common Ancestor (LECA) (Azimzadeh, 2021, 2014; Azimzadeh and Marshall, 2010; Carvalho-Santos et al., 2010; Hodges et al., 2010). Over millions of years of evolution, the molecular architecture of the centriole has been preserved in parts in many species, but disappeared in some cases concomitantly with the loss of motile flagellum (Azimzadeh and Marshall, 2010), such as in yeasts, amoebozoa and flowering plants (Nabais et al., 2020). Nevertheless, yeasts conserved a rudimentary organelle, the spindle pole body (SPB), which carries some similarities with the centriole, such as duplication and assembly process that are tightly linked to the cell cycle (Seybold and Schiebel, 2013).

In yeast, the SPB duplication is characterized by the formation of a half bridge structure, made of Cdc31 and Sfi1, that provides a platform for SPB duplication (Bouhlel et al., 2015; Jaspersen et al., 2002; Kilmartin, 2003; Spang et al., 1995). Intriguingly, SFI1 and centrins are also present in mammalian centrosomes but whether they form a complex at centrioles and are involved in centriole duplication remained unclear. In this paper, we establish that SFI1 is a *bona fide* centriolar protein that is recruited early during centriole biogenesis at its growing distal end. Importantly, we found that a pool of centrins display a similar localization at the distal end of centrioles in addition to the previously described inner scaffold localization (Le Guennec et al., 2020; Steib et al., 2020). In addition, we noticed that the Centrin3 signal is slightly less extended (**Figure 1M**) but this difference might possibly be due to either the quality of the antibody or reflecting a real difference between the two centrins. By measuring at nanometric scale precision the distance between centrins and SFI1 signals, we found that these proteins are about 20 nm distant from each other, which is negligible if we consider that the size of 15 SFI1 repeats is about 60 nm long (Li et al., 2006), and that SFI1 antibody recognizes SFI1 C-terminus and not the repeats region. Moreover, biophysical data showed that these proteins interact directly *in vitro* (Martinez-Sanz et al., 2010, 2006). Taken together, we can assert that the complex SFI1/centrin is conserved in mammals and that it is localized at the distal end of centrioles, from the early stages of procentriole assembly. Additionally, we demonstrated that SFI1 is critical for centrin targeting at the distal end of centrioles. On the other hand, we established that POC5 drives the localization of centrin at the central core region of centrioles while it does not affect the distal pool of centrin nor SFI1. Altogether, these results highlight the presence of two distinct complexes containing centrin: one at the distal end of the centriole dependent on SFI1 and one at the central core relying on POC5.

In SFI1 depleted cells, we observed a decrease in centrin signal that is consistent with the reduced GFP-centrin1 levels seen in a screen for centriole biogenesis factors (Balestra et al., 2013). However, in contradiction with previous reports (Balestra et al., 2013; Kodani et al., 2019), we did not observe defects in centriole duplication in U2OS cells as well as a decrease in CP110 recruitment at centrioles, neither by standard immunofluorescence procedure nor by expansion microscopy. These observations indicate that the SFI1/centrin complex is present at centrioles but may not participate in the initiation of centriole duplication in humans, unlike its yeast counterpart. Instead, SFI1 depletion leads to structurally abnormal centrioles, which lose their canonical organization. Therefore, we suggest that the SFI1/centrin complex may serve as procentriole growth guidance from its tip or could possibly also play a role in the maintenance of centriole architecture once growth is completed. In addition, we also unveiled that SFI1 is important for ciliogenesis but it remains to be determined whether this is due to the observed structural defects upon SFI1 depletion.

Another intriguing question concerns the molecular organization of SFI1/centrins inside centrioles. In yeast, Sfi1 molecules organize into two anti-parallel arrays connected by Sfi1 C-termini, with the N-termini oriented towards the SPB cores (Li et al., 2006). In our study, we focused on localizing the C-terminus of human SFI1. If the C-terminal interactions are conserved in spite of strong sequence divergence between yeast and human SFI1 outside of centrin-binding domains, we could imagine that SFI1 C-termini could interact in the center of the lumen, while N-termini domains would radially extend towards the periphery facing centriolar microtubule walls at the centriole’s distal end, similar to the cartwheel structure found in the proximal region. Such a radial structure has never been observed in the centriole, but it is likely that another type of assembly exists there. Indeed a recent study has shown that C2CD3 and LRRCC1 also localize at the luminal distal end of the human centriole (Gaudin et al., 2021), similarly to SFI1 and centrin, and delineate a structure reminiscent of the acorn, a filamentous density observed by electron microscopy in pro- and mature *Chlamydomonas* basal bodies (Gaudin et al., 2021; Geimer, 2004). In addition, the acorn is accompanied by a V-shaped filament system that has been proposed to be composed of centrin (Geimer and Melkonian, 2005). It is therefore possible that SFI1 and centrin are part of this V-shaped filament system and associate with other distal extremity centriolar proteins, such as C2CD3 and LRRCC1, to ensure a proper centriole formation.

It will also be necessary to better understand the function of the different centrins in human and whether they form separate complexes with SFI1 or can co-assemble in the same ones if co-expressed in the same cell. Since Centrin-1 is only expressed in the testis and Centrin-4 in ciliated cells and in retina, they might assume cilia/flagella specific functions while Centrin-2 or Centrin-3 could be mainly involved in centriole biogenesis. Solving these questions would certainly help fully understand the function of SFI1/centrin complex in guiding centriole biogenesis in human cells.

By identifying a new mechanism guiding centriole growth and regulating centriole architecture depending on the evolutionary conserved proteins SFI1 and centrins, our work may also bring new insights on the evolution of centrosome biogenesis. In the absence of a better knowledge of the molecular mechanisms by which the SFI1/centrin complex achieves its potential role in guiding centriole growth, it is difficult to imagine what common principles it may share with the SPB duplication process in yeast, although a structural role in holding duplicated SPBs or centriole walls together could be envisaged. On the other hand, Ecdysozoa such as flies and worms contain the evolutionary conserved proteins SAS-6, SAS-5/ANA2/STIL, SAS-4/CPAP, CEP135/Bld10p and POC1 critical for duplication and assembly of the procentriole but lack SFI1 and centrin genes (Ito and Bettencourt-Dias, 2018; Kilmartin, 2003). Remarkably, they also display atypically short centrioles where the guiding function of the SFI1/centrin complex could then be dispensable. In this clade, alternative guiding mechanisms might allow axoneme growth in the small subset of cells that do assemble cilia or flagella.

## Acknowledgements

We thank Nikolai Klena and Olivier Mercey for critical reading of the manuscript. We thank Paul Conduit for support to I.B.B. The authors greatly acknowledge the Cell and Tissue Imaging (PICT-IBiSA), Institut Curie, member of the French National Research Infrastructure France-BioImaging (ANR10-INBS-04), the Nikon Imaging Centre at Institut Curie-CNRS as well as the Bioimaging Center at Unige (Geneva, Switzerland). This work is supported by a grant PJA 20151203291 from Fondation ARC pour la Recherche sur le Cancer attributed to Anne Paoletti, the Swiss National Science Foundation (SNSF) PP00P3_187198 and by the European research Council ERC ACCENT StG 715289 attributed to Paul Guichard as well as by a grant PJA3 2020060002055 from Fondation ARC pour la Recherche sur le Cancer to JA. Anne Paoletti is a member of Labex CelTysPhyBio (ANR-11-LABX-0038) and Cell(n)Scale (ANR-10-IDEX-0001-02). Imene B. Bouhlel received doctoral fellowships from Université Paris-Sud and a 4^th^ year PhD fellowship from Fondation ARC pour la Recherche sur le Cancer and a travel fellowship from Labex CelTysPhyBio. Eloïse Bertiaux received an EMBO fellowship ALTF 284-2019.

## Author contributions

I.B.B. and M.H.L performed and analyzed the experiments of the paper with the help of E.B., A.G. and S.B.. A.P, V.H. and P.G conceived, supervised and designed the project. I.B.B, M.H.L, A.P, V.H. and P.G wrote the manuscript with the input from all authors. J.A. generated the SFI1 homemade antibody in the laboratory of M.B. M.B. provided expertise all along the project.

## Material and Methods

### Human cell lines and Cell Culture

RPE-1 cells were cultured in DMEM-F12 medium supplemented with 10% fetal calf serum and 1% Penicillin-Streptomycin at 37°C and 5% CO2. To induce ciliogenesis, cells were starved from serum during 48h. U2OS cells were cultured in DMEM supplemented with GlutaMAX (Life Technology), 10% tetracycline-negative fetal calf serum (life technology), penicillin and streptomycin (100 µg/ml) at 37°C and 5% CO2. Cells were tested for mycoplasma contaminations regularly.

### SFI1 depletion using siRNAs

The ON-TARGET plus Non-targeting Pool siRNA was used as control and purchased from Dharmacon (Catalogue #D-001810-10-20). It is composed of a pool of four siRNAs designed and microarray tested for minimal targeting of human. The siRNAs used to deplete SFI1 were designed as described in (Balestra et al., 2013) and purchased from Eurogentec. The sequences are as follows: siSfi1#1 (AAGCAAGTACTCATTACAGAA-dTdT) and siSfi1#2 (AAGGTTGTCTCTGCAGTGAAA-dTdT). Silencer select negative control siRNA1 were purchased from Thermo Fisher (4390843, Thermo Fisher). RPE-1 cells were plated (50 000 cells) on 12 or 15 mm coverslips in a 6-well plate and 10 nM siRNA-1 or -2 was transfected using RNAi MAX reagents (Invitrogen) according the manufacturer protocol. Cells were analyzed 72 hours after transfection. Since both siRNAs gave similar results (data not shown), we concentrated on siRNA#1 in the U-ExM experiment.

### POC5 depletion using siRNAs

U2OS cells were plated onto coverslips in a 6-well plate at 100 000 cells/well 24 hours prior transfection. Cells were transfected either with 50 nM silencer select negative control siRNA1 (4390843, Thermo Fisher) or 25 nM siPOC5 (sequence Sense siPOC5-1: 5’ CAACAAAUUCUAGUCAUACUU 3’ and antisense: 5’ GUAUGACUAGAAUUUGUUGCU 3’, adapted from (Azimzadeh et al., 2009) using Lipofectamine RNAimax (Thermo Fischer Scientific). Medium was changed 6 hours post-transfection and cells were analyzed 72 hours post-transfection. Note that POC5 depletion was partial (Figure S2O).

### Antibodies

#### SFI1 Antibody purification

The SFI1 antibody was raised in rabbit against the GST-fused C-terminal domain of SFI1 (aa1021 to aa1240) (Figure S1) and affinity-purified on AminoLink® Coupling Resin (20381 Thermo Fisher) coupled to the MBP-fused C-terminal domain (using the same aa sequence).

The antibodies used in this study are the following: SFI1 (13550-1-AP, Proteintech Europe), home-made SFI1 (this study, 1:200), γ-tubulin (sc-7396, Santa Cruz Biotechnology, Inc., 1:500), Centrin (clone 20H5, 04-1624, Millipore, 1:500), Centrin3 (H00001070-M01, Abnova), PCNA (mAb #2586, Cell Signalling Technology, 1:1000), HsSAS-6 (sc-81431, sc-98506 Santa Cruz Biotechnology, Inc.), STIL (A302-441A-T, Bethyl), CP110 (EPP11816, Elabscience, 1:500), Acetylated tubulin (Institut Curie Recombinant antibodies Platform, 1:75), β-tubulin (AA344, scFv-S11B) and α-tubulin (AA345, scFv-F2C) (Nizak et al., 2003), α-tubulin (1:500, Abcam, ab18251) were purchased from the indicated suppliers. Primary antibodies were used at the concentration mentioned above when used for classical immunofluorescence and at 1:250 for U-ExM experiments. Secondary fluorescent antibodies were purchased from Invitrogen (A11008, A11004, A11029 and A11036, Invitrogen, ThermoFisher) and used at 1:800 dilutions for standard immunofluorescence experiments and 1:400 for U-ExM.

### Immunofluorescence microscopy

For immunofluorescence, cells were grown on 12 mm coverslips and fixed at -20°C with cold methanol for 3 min. Fixed cells were then incubated with the primary antibodies for 1h at room temperature, washed with PBS and subsequently incubated with the secondary antibodies conjugated with Alexa Fluor-488, 594 or 647. DNA was counterstained with DAPI solution. Samples were mounted in Mowiol and observed with a fluorescence microscope (Upright Leica DMI-5000B) equipped with a CCD Camera 1392×1040 (CoolSnap HQ2 pixel: 6.45 µm from Photometrics). Images were acquired and processed using Metamorph software (Molecular Devices). For the quantification of fluorescence intensity (Figure 3B), maximal projections were used using Fiji (Schindelin et al., 2012). Confocal centriolar intensities were assessed by individual plot profil along a linescan of 30 pixel on each pair of mature centrioles. For each experiment, all values were normalized on the average value of the control cells to obtain the relative intensity (A.U.). An average of all normalized measures was generated and plotted in GraphPadPrism7.

### Ultrastructure Expansion Microscopy (U-ExM)

The following reagents were used in U-ExM experiments: formaldehyde (FA, 36.5– 38%, F8775, SIGMA), acrylamide (AA, 40%, A4058, SIGMA), N, N’-methylenbisacrylamide (BIS, 2%, M1533, SIGMA), sodium acrylate (SA, 97–99%, 408220, SIGMA), ammonium persulfate (APS, 17874, Ther-moFisher), tetramethylethylendiamine (TEMED, 17919, ThermoFisher), nuclease-free water (AM9937, Ambion-ThermoFisher) and poly-D-Lysine (A3890401, Gibco).

RPE-1 and U2OS cells were grown on 12 mm coverslips and processed for expansion as previously described (Le Guennec et al., 2020; Steib et al., 2020). Briefly, coverslips were incubated in 2% AA + 1.4% FA diluted in PBS for 5 hr at 37°C prior to gelation in monomer solution (19% sodium acrylate, 0.1% BIS, 10% acrylamide) supplemented with TEMED and APS (final concentration of 0.5%) for 1 hr at 37°C. Denaturation was performed for 1h30 at 95 °C and gels were stained as described above. For each gel, a caliper was used to accurately measure its expanded size. The gel expansion factor was obtained by dividing the size after expansion by 12 mm, which corresponds to the size of the coverslips use for sample seeding. Measurements of lengths and diameters were scaled according to the expansion factor of each gel.

### Image analysis

Expanded gels were mounted onto 24 mm coverslips coated with poly-D-lysine (0.1 mg/ml) and imaged with an inverted Leica TCS SP8 microscope or using a 63 × 1.4 NA oil objective with Lightening mode at max resolution, adaptive as ‘Strategy’ and water as ‘Mounting medium’ to generate deconvolved images. 3D stacks were acquired with 0.12 µm z-intervals and an x, y pixel size of 35 nm. Length coverage and diameter quantification was performed as previously published in (Le Guennec et al., 2020). For the measurement of SFI1 intensity (Figure S2D), the Fiji plot profile tool was used to obtain the fluorescence intensity profile from proximal to distal for tubulin and of SFI1 from the same line scan. Roundness was calculated on perfectly imaged top views of centrioles by connecting tubulin peaks on ImageJ. To generate the panels in Figure 1E, H, K and L, we used two homemade plugins for ImageJ as described previously (Borgne et al., 2021).

### Statistical analysis

The comparison of two groups was performed using an unpaired two-sided Student’s t-test or its non-parametric correspondent, the Mann-Whitney test, if normality was not granted because rejected by Pearson test. The comparisons of more than two groups were made using one-way ANOVAs followed by post-hoc tests as indicated in corresponding figure legend to identify all the significant group differences. N indicates independent biological replicates from distinct samples. Every experiment was performed at least three times independently on different biological samples unless specified. No statistical method was used to estimate sample size. Data are all represented as scatter dot plot with centerline as mean, except for percentage quantifications, which are represented as histogram bars. The graphs with error bars indicate SD (+/-) and the significance level is denoted as usual (*p<0.05, **p<0.01, ***p<0.001, ****p<0.0001). All the statistical analyses were performed using Excel or Prism7 (Graphpad version 7.0a, April 2, 2016).

### Contact for reagent and resource sharing

Further information and requests for resources and reagents should be directed to Anne Paoletti, Paul Guichard, and Virginie Hamel.

## Figures Legends

**Figure S1.**
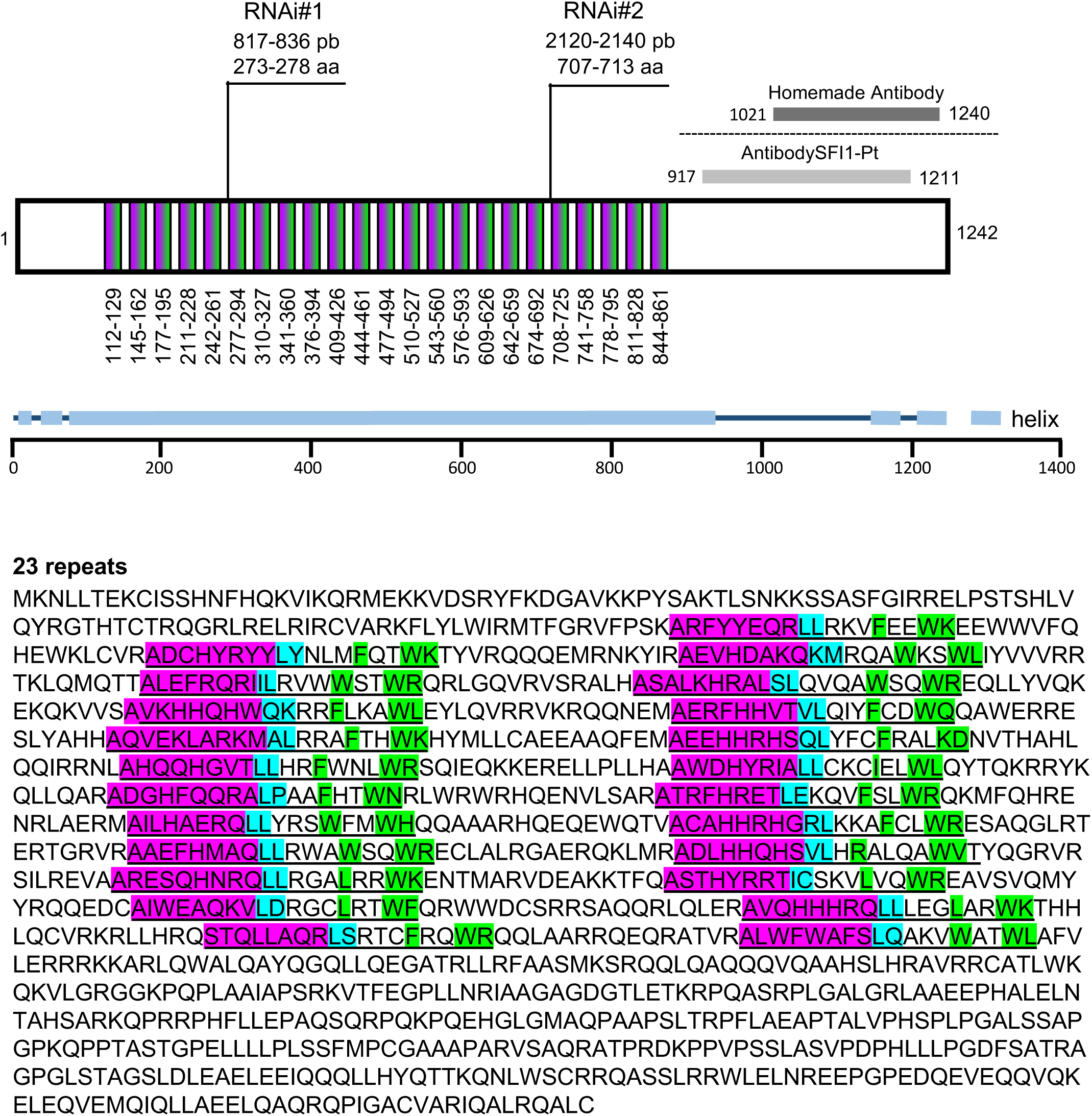
Human SFI1 protein. Schematic representation of the 1242 amino acid human SFI1 protein. Note the relative position of the two antibodies used in this study (home-made antibody and SFI1-Pt, Proteintech) as well as the regions targeted by the two siRNAs used in this study. As both siRNA gave similar results, only the siRNA1 was used throughout the manuscript. The 23 SFI1 repeats are underlined. The colors magenta, cyan and green highlight the SFI1 motif.

**Figure S2.**
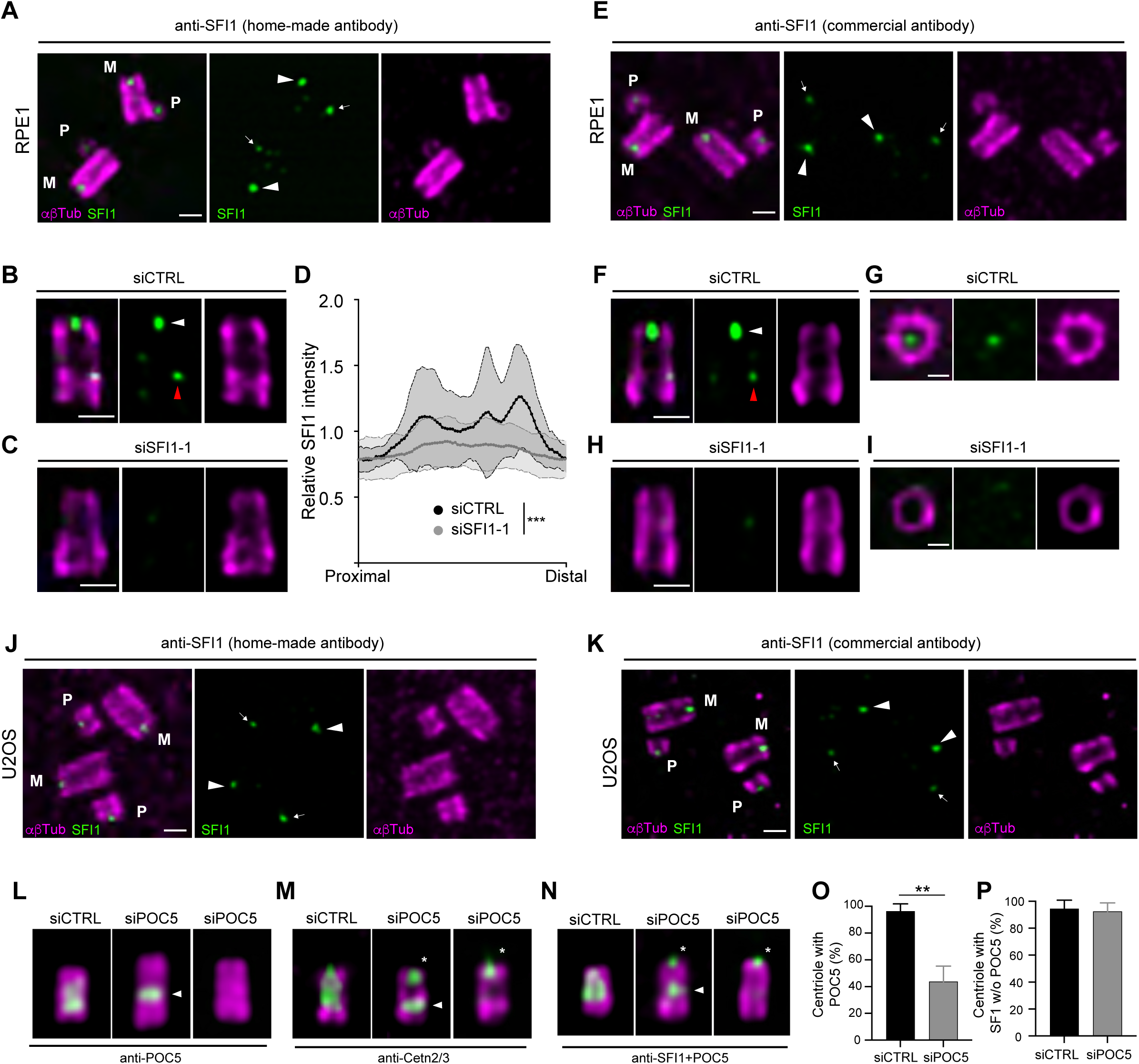
SFI1 localization at centrioles revealed by expansion microscopy. **(A)** Representative confocal image of expanded RPE-1 centrioles stained for α/β-tubulin (αβTub, magenta) and SFI1 (green, home-made antibody). Arrowheads point to SFI1 signal at mature centrioles while the thin arrows highlight SFI1 signal at procentrioles. M stands for mature centrioles and P for procentriole. Scale bar: 200 nm. (**B, C**) Representative confocal image of expanded RPE-1 centrioles treated with siControl (**B**) or siSFI1-1 (**C**), stained with α/β-tubulin (αβTub, magenta) and SFI1 (green, home-made antibody). White arrowheads point to SFI1 signal at the distal end of centrioles while red arrowhead points to a faint proximal signal. Scale bar: 200 nm. **(D)** Relative SFI1 intensity in the indicated conditions showing a significant decrease in siSFI1-1-treated cells. Averages of AUC (area under the curve) +/- SD are as follows: siCTRL: 0.79 +/- 0.2, siSFI1-1: 0.55 +/- 0.3. N=24 for siCTRL and 50 for siSFI1-1 from 4 independent experiments. Mann-Whitney test, p<0.0001. **(E)** Representative confocal image of expanded RPE-1 centrioles stained for α/β-tubulin (αβTub, magenta) and SFI1 (green, commercial antibody 13550-1-AP). M stands for mature centriole and P for procentriole. White arrowheads indicate SFI1 signal at mature centrioles and thin white arrows, SFI1 signal at procentrioles. Scale bar: 200 nm. **(F-I)** Representative confocal image of expanded RPE-1 centrioles treated with siCTRL (**F, G**) or siSFI1-1 (**H, I**), stained with α/β-tubulin (αβTub, magenta) and SFI1 (green, commercial antibody 13550-1-AP). White arrowheads point to SFI1 signal at the distal end of centrioles while red arrowhead points to a faint proximal signal. **(J, K)** Representative confocal image of expanded U2OS centrioles stained with α/β-tubulin (αβTub, magenta) and SFI1 (green, home-made antibody (**J**) or commercial antibody (**K**)). White arrowheads point to SFI1 signal at mature centrioles and white arrows highlight SFI1 signal at procentrioles. M stands for mature centrioles and P for Procentrioles. Scale bar: 200 nm. **(L-N)** Representative widefield images of expanded U2OS centrioles treated with siCTRL or siPOC5, stained with α/β-tubulin (αβTub, magenta) and POC5 (green, **L**), Cetn2/3 (green, **M**) or SFI+POC5 (green, **N**). White arrowheads indicate the remaining proximal belt of POC5 sometimes observable in siPOC5 treated cell when depletion is incomplete (**L**, middle panel). Note that centrin behavior seems to follow POC5 upon POC5 depletion (**M**, middle panel). Asterisks indicate the presence of the distal dot of centrin and SFI1 in POC5-depleted cells. Scale bar: 200 nm. (**O**). Quantification of the siPOC5 efficiency at centrosomes. Average +/- SD are as follows: siCTRL= 96.4 % +/- 5.4; siPOC5= 43.8%+/- 11.6, indicating that on average a bit more than half of centrioles are depleted for POC5 under these experimental conditions. N=5 independent experiments (100 cells per experiment), **p value=0.002, Mann-Whitney test. (**P**). Percentage of depleted centrioles (without POC5 staining) containing SFI1 as a distal dot in siCTRL and siPOC5 treated cells. Average +/- SD are as follows: siCTRL= 94.5 % +/- 6.4; siPOC5= 92.5%+/- 6.4, demonstrating that SFI1 localization is not impacted by POC5 depletion under these experimental conditions. N=2 independent experiments (100 cells per experiment).

**Figure S3.**
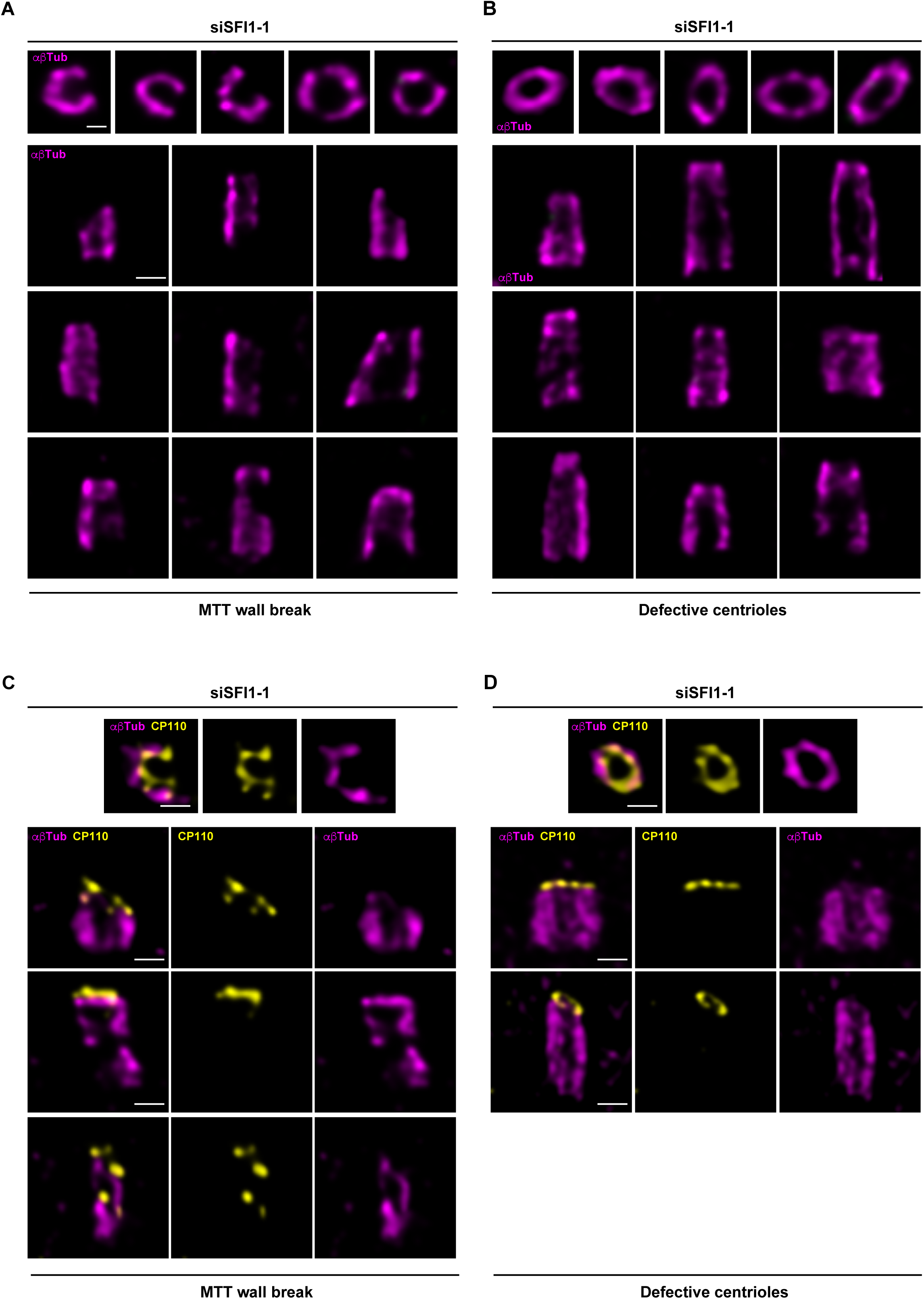
Gallery of defective centrioles in SFI1-depleted RPE-1 cells. (**A, B**) Confocal images of expanded centrioles from SFI1-depeted RPE-1 stained for α/β-tubulin (Magenta). Top view (top panel) and side view (bottom panels) of broken centriole (**A**) and abnormal but not broken (**B**) stained for (αβTub, magenta) and SFI1 (green). Scale bar: 200 nm. (**C, D**) Confocal images of expanded centrioles from SFI1-depeted RPE-1 stained for α/β-tubulin (Magenta) and CP110 (yellow). Top view (top panel) and side view (bottom panels) of broken centriole (**C**) and abnormal but not broken (**D**) stained for (αβTub, magenta) and CP110 (yellow). Scale bar: 200 nm.

**Video 1. U-ExM expanded control and SFI1-depleted centrioles**.

U-ExM expanded centriole from RPE-1 cell treated with scramble or siSFI1 siRNA and stained for α/β-tubulin (Magenta) and SFI1 (Green, homemade antibody). Z-stack acquired every 0.14 µm from the proximal to distal end of the centriole. Note the loss of the characteristic roundness of the centriole in the SFI1-depleted centriole.

